# *Afrothismia lovettii* sp. nov. (Afrothismiaceae) Critically Endangered (Possibly Extinct) and endemic to the Kihansi Falls Forest, Tanzania

**DOI:** 10.64898/2026.06.02.729641

**Authors:** Martin Cheek, Jon C. Lovett

**Affiliations:** Herbarium, Royal Botanic Gardens, Kew, Richmond, Surrey, TW9 3AE, UK; School of Geography, University of Leeds, Leeds LS2 9JT, UK

**Keywords:** achlorophyllous mycotroph, conservation, Eastern Arc, *Glomales*, Glomeromycota, *Kihansia*, *Kupea*, Udzungwa Mts, World Bank

## Abstract

The fully mycotrophic (or mycoheterotrophic) non-photosynthetic *Afrothismia lovettii* Cheek sp. nov. (Afrothismiaceae), is described and illustrated from the Kihansi Gorge Forest of the Udzungwa Mts, Tanzania where a World Bank-funded hydroproject may have resulted in its extinction, together with the possible extinction of two other narrowly endemic species of mycoheterophic plant, *Kihansi*a *lovettii* and *Kupea jonii* (Triuridaceae). The co-incidence of these three species makes (or made) the Kihansi Falls the most important site for species diversity of mycotrophic plants in eastern Africa. In terms of numbers (three) of strict endemic species of this lifeform, the Kihansi Falls is or was, the most important site in Africa together with Mt Kupe in Cameroon. *Afrothismia lovettii* is similar and may be related to *Afrothismia mhoroana* Cheek of the Uluguru Mts with which it was confused, differing inter alia in the mainly white, short (6.5–10 mm long), ±monomorphic perianth lobes (vs red, 12 and 20 mm long, di- or tri-morphic) and glabrous (vs hairy) adnate staminal filaments. The provisional extinction risk assessment for *Afrothismia lovettii* is Critically Endangered (Possibly Extinct) (CR B1ab (iii)+2ab(iii)) using the IUCN 2012 categories and criteria, due to the small area of occupancy and the forest habitat degradation (loss of waterfall spray) resulting from the construction of the World Bank financed Kihansi dam.

## Introduction

Discovery of a specimen image (P.G. Hawkes in *Neyland* 1936, LSU) on gbif.org of an unknown *Afrothismia* from Kihansi Falls, Udzungwa Mts of Tanzania prompted re-examination of *Lovett* 5063 (DSM, K), a specimen also from this site previously tentatively identified as *A. cf. winkleri* by the first author (Cheek 2004a), later misidentified as *A. mhoroana* Cheek, an endemic of the Uluguru Mts, also in Tanzania. The re-examination was supported by the newly available photos (Fig. 1 & 2) from the second author and collector of *Lovett* 5063, and also by the availability of previously overlooked duplicates. Although *Lovett* 5063 and *Neyland* 1936 (Fig. 3) have similarly sized, similarly shaped and decorated, and similarly coloured flowers to *A. mhoroana*, there are so many points of difference (Table 1) that there is no doubt that they represent a separate species, here characterised and formally named as *Afrothismia lovettii* Cheek.

**Table 1.** Diagnostic characters separating *Afrothismia lovettii* from *A. mhoroana*. Data on the last species from Cheek & Jannerup (2006).

|  | *Afrothismia mhoriana* | *Afrothismia lovettii* |
| --- | --- | --- |
| Perianth lobes | Dimorphic (or trimorphic);<br>c. 20 mm (dorsal lobes) and | Monomorphic (lobes<br>subequal in length) 6.5–10<br>mm long. |
|  | c. 12 mm (ventral lobes) long. |  |
| Indumentum of adnate staminal filaments | Densely hairy | Glabrous |
| Perianth lobe colour | Red | White (sometimes yellow at base) |
| Corona (perianth mouth) shape | Spreading (as in the mouth of a trumpet) | Shortly cylindrical |
| Anther connective (dorsal view) | Wider than anther cells, concealing them completely | Narrower than the anther cells which are visible. |
| Geographic range | Uluguru Mts | Udzungwa Mts |

**Fig 1.**
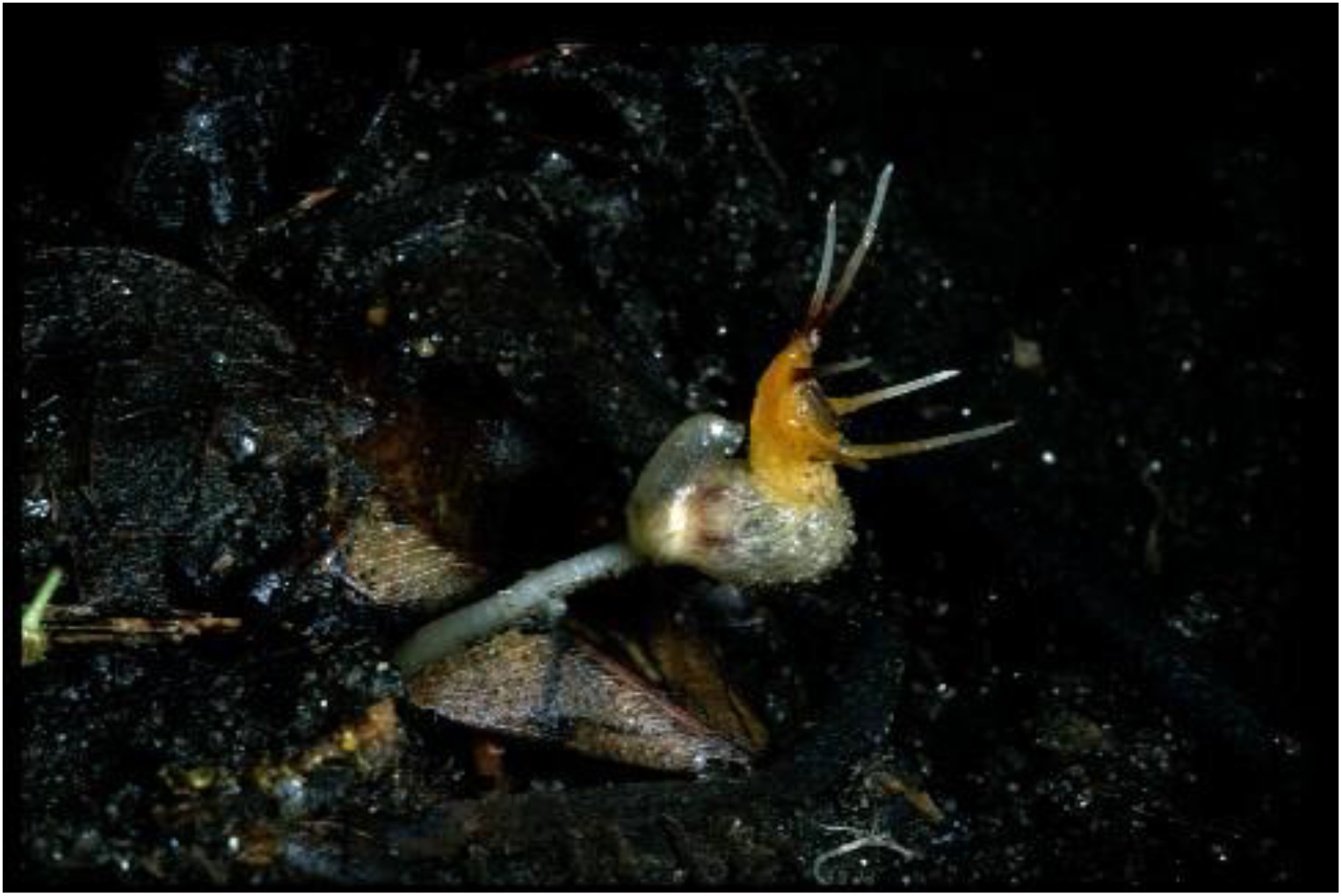
Afrothismia lovettii. Photo of a plant of the type specimen *Lovett* 5063, side view of flower, taken in situ at Kihansi Gorge in 1997. Photo by Jon Lovett.

**Fig 2.**
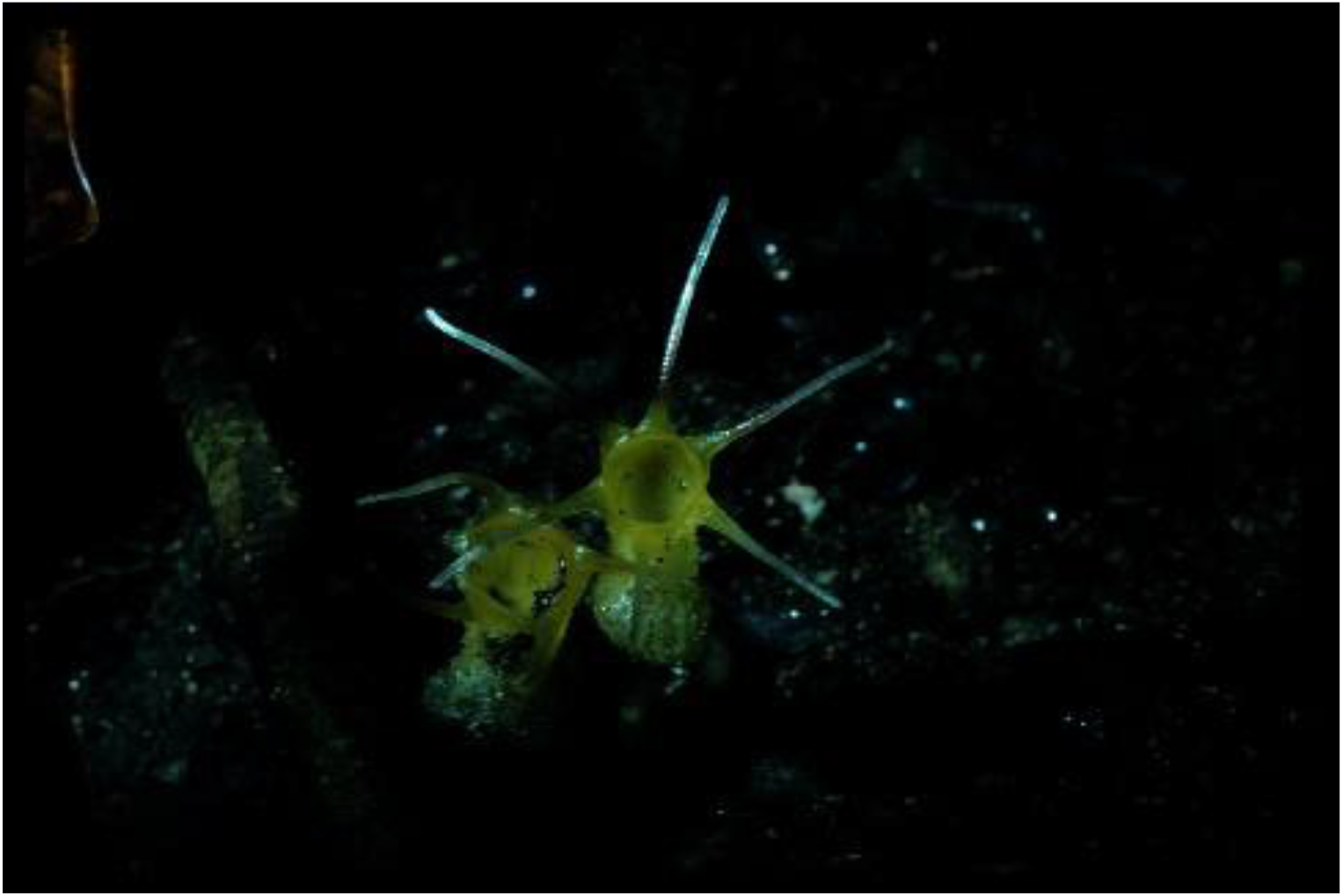
Afrothismia lovettii. Photo of a plant of the type specimen *Lovett* 5063, view of mouth of perianth tube, note the equal perianth lobes. Photo by Jon Lovett.

**Fig 3.**
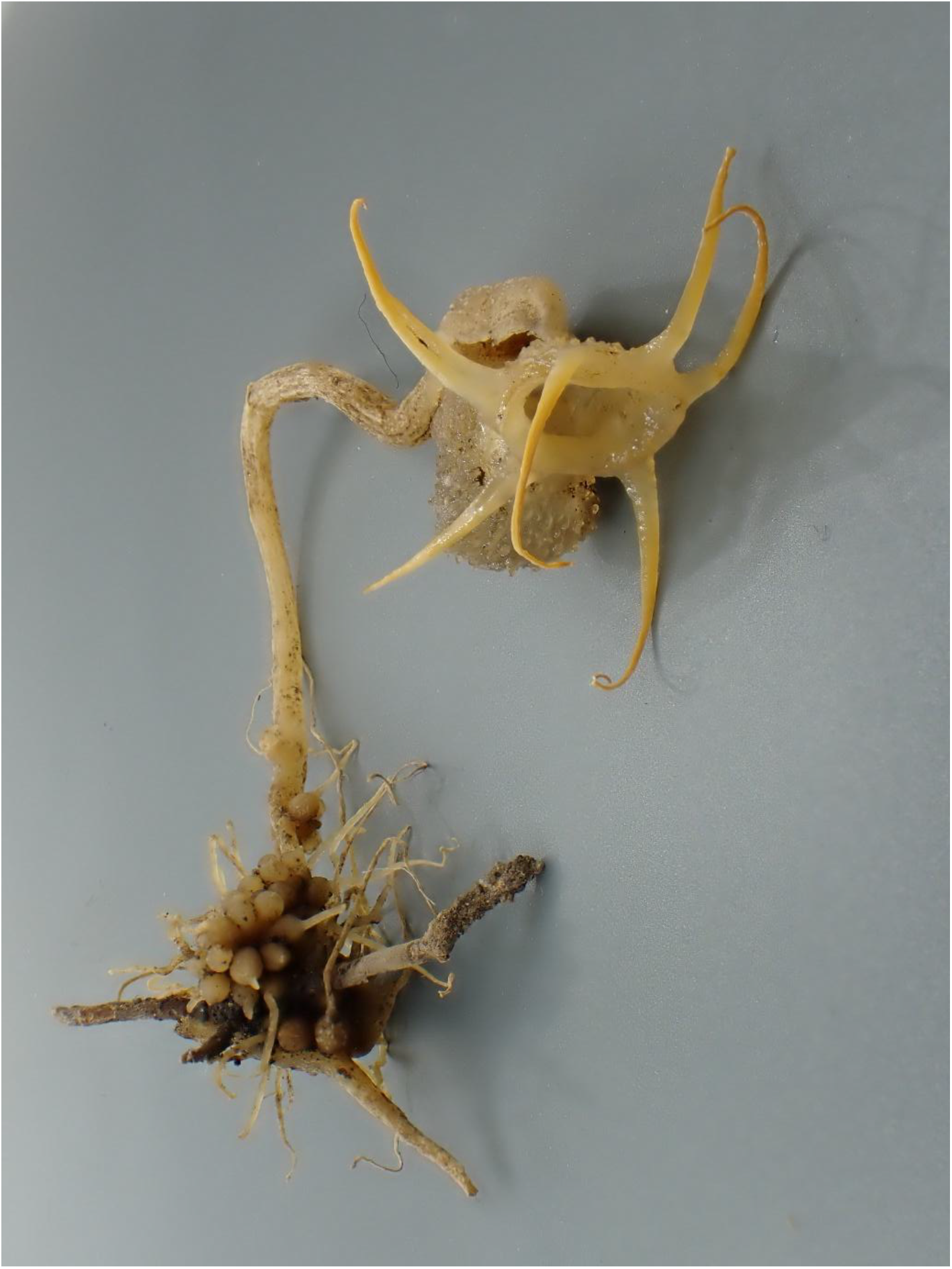
Afrothismia lovettii. Photo of liquid-preserved plant of Hawkes in *Neyland* 1936, note the horizontal stem and bulbil clusters at base of vertical stem. Photo by Peter Hawkes

**Fig 4.**
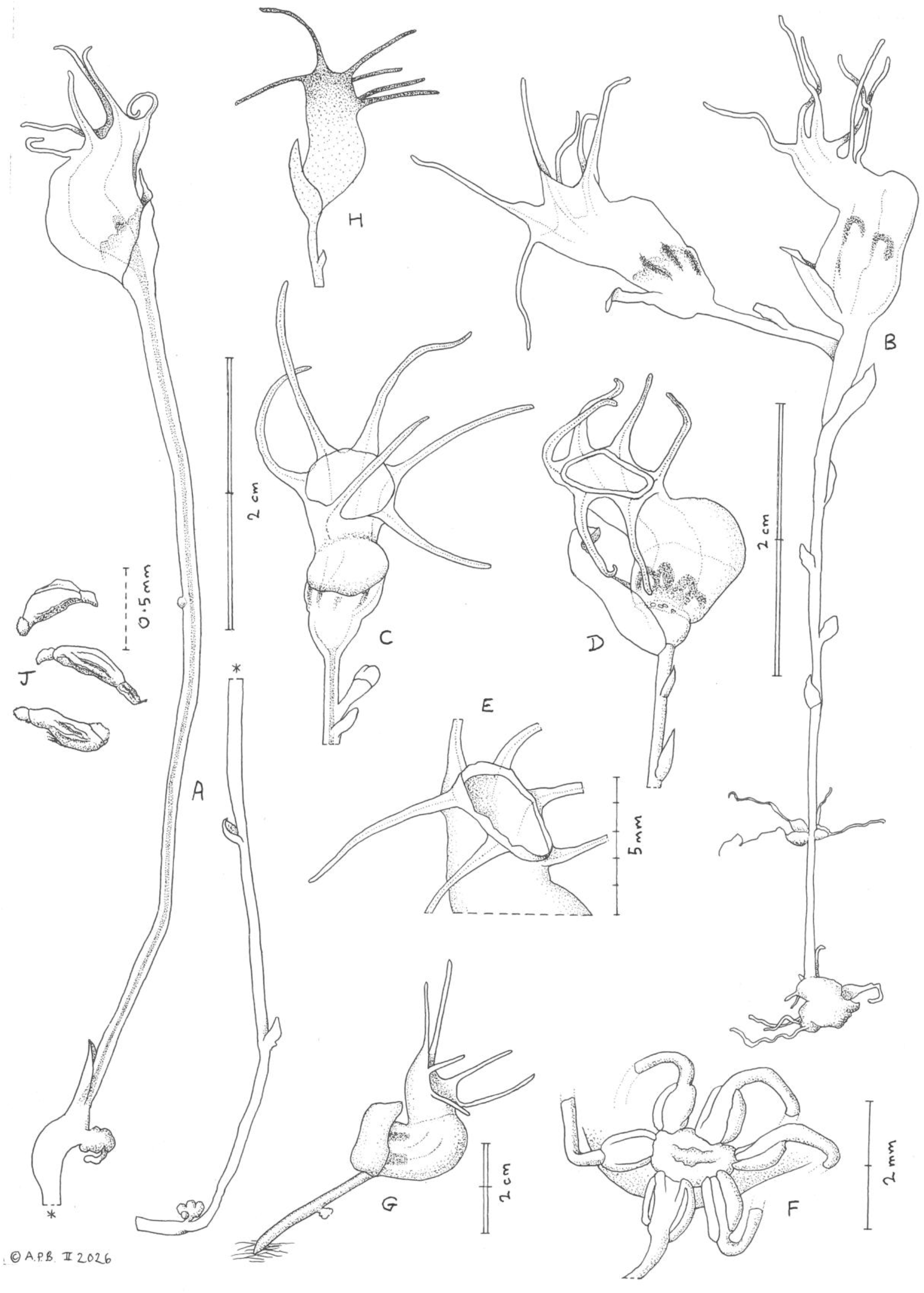
Afrothismia lovettii. A flower and stem (in two parts); B ibid with root bulbil clusters attached; C ventral view of flower; D hydrated flower, side view, androecium shown in silhouette; E mouth of corolla tube showing short corona (hydrated flower); F view from above of stamens attached to the stigma (hydrated flower); G live plant in situ (from photo); H outline of flower showing distribution of pigment (darker areas = yellow); J seeds, side view. All drawn from *Lovett* 5063 by Andrew Brown.

Fully mycoheterotrophic non-photosynthetic plant species, often known as achlorophyllous mycotrophic plants, mycoheterophs (Merckx 2013) or saprophytes, are remarkable for lacking all chlorophyll and being completely dependent on vascular arbuscular mycorrhizal fungi for sustenance. For reviews of the species diversity, species assemblages, ecology and geography of the African plant families and genera with this life form see Cheek & Ndam (1996), Cheek & Williams (1999), Cheek (2006). The genus *Rhizophagus* P.A. Dang (Glomerales, Glomeromycota) has been implicated as the fungal partner of *Afrothismia* (Franke et al. 2006).

Afrothismiaceae Cheek & Soto Gomez (Cheek et al. 2023), with a single genus, *Afrothismia*, are confined to tropical continental Africa. *Afrothismia* differs from the genera of Thismiaceae by the circumscissile fruit dehiscence, axile placentation, elevation of the placental mass from the fruit by a placentophore, annulus inserted deep inside the perianth tube; stamens inserted below the annulus; anthers usually adnate to stigma and rhizomes with clusters of usually spherical tubers among other characters (Cheek et al. 2023). For further information on the phylogeny, anatomy and morphological diversity of the family see Cheek et al. (2023). Eighteen species have been formally described or are in the course of publication (now 19, with this paper), but there appear to be seven further undescribed species in Cameroon and Gabon (Cheek et al. 2023).

*Afrothismia* was erected by Schlechter (1906), based on *Thismia winkleri* Engl. (Engler 1905) with a second species, *A. pachyantha* Schltr., both from Cameroon. With *A. insignis* Cowley (Tanzania) and Ugandan *Afrothismia winkleri* var. *budongensis* Cowley (now *A. ugandensis* Cheek, Cheek et al. 2024) the range of the genus was extended to E Africa (Cowley 1988).

With the publication of the Cameroonian species, *A. gesnerioides* H.Maas (Maas-van der Kamer 2003) the original two species known for the genus had increased to four in a century. In the following 15 years, the number of formally described new species quadrupled to 16, with discoveries from Gabon (Dauby et al. 2008), Kenya (Cheek 2004b), Malawi (Cheek 2009) and Tanzania (Cheek & Jannerup 2006). The largest number of discoveries in the period was made in Cameroon. Here, eight new species of *Afrothismia* have been published, one extending to Nigeria (Franke 2004; Franke et al. 2004; Sainge & Franke 2005; Sainge et al. 2005; Cheek 2007; Sainge et al. 2013; Cheek et al. 2019). Most of the Cameroonian species fall within the Cross-Sanaga interval (Cheek *et al*. 2001), which holds the highest flowering plant species (Barthlott et al. 1996) and generic (Dagallier et al. 2020) diversity per degree square in Tropical Africa. In contrast, only two specimens were listed for Gabon, neither identified to species (Sosef et al. 2006). Eight of the Cameroonian *Afrothismia* species feature in the Red Data Book of Cameroon Plants (Onana & Cheek 2011) and most species of the genus are Critically Endangered (Cheek et al. 2023). One Cameroonian species is considered extinct after targeted searches (Cheek et al. 2019), while several others have not been recorded alive in several decades and so may also be extinct, e.g. *A. zambesiaca* Cheek (Cheek 2009).

Although species niche modelling (Sainge et al. 2017) had indicated that *Afrothismia* might occur in West Africa, west of the Dahomey gap, the areas predicted were Liberia, Cote D’Ivoire and Sierra Leone, not in Guinea where it was recently reported (Cheek et al. 2026).

## Materials & methods

Names of species and authors follow IPNI (continuously updated). Herbarium material was examined with a Leica Wild M8 dissecting binocular microscope fitted with an eyepiece graticule measuring in units of 0.025 mm at maximum magnification. The drawing was made with the same equipment with a Leica 308700 camera lucida attachment. Herbarium material was softened with soapy water to rehydrate it before dissection as per the Herbarium Handbook (Davies et al. 2023). The following herbaria were inspected for specimens: B, BM, EA, HNG, K, LSU, MHU, SRGH, YA and WAG. Herbarium codes follow Index Herbariorum (Thiers, continuously updated).

The use of technical terms follows Beentje & Cheek (2003) and the format of the description follows those in other papers describing new species in *Afrothismia*, e.g. Cheek et al. (2026). The specimens cited have all been seen by one or other of the authors. The conservation assessment follows the IUCN (2012; 2024) Red List of Threatened Species categories and criteria, and Red List Guidance, respectively.

## Results Taxonomy

### Afrothismia lovettii *Cheek* sp nov

Type: “Tanzania, Southern Udzungwa Escarpment, 8° 37′S, 35°51′E” (erroneous), 770 m, fl.fr. 21 March 1997, *J. Lovett* 5063 (holo-K[K000593397]; iso-K 000541643, iso-DSM n.v.). Figs 1–4 Diagnosis: differing from *Afrothismia mhoroana* Cheek in the perianth lobes (tepals) mainly white, short, ±monomorphic 6.5–10 mm long (vs red, di or tri-morphic 12 and 20 mm long); glabrous (vs hairy) adnate staminal filaments; perianth mouth with corona shortly cylindric (vs spreading like a trumpet). See additional diagnostic characters in Table 1.

#### Description

*Non-photosynthetic perennial mycoheterotrophic herb* to 4 cm tall (field notes), only the flower and fruit and a few cm of stem emerging above the leaf-litter. *Stem* (rhizome), succulent, translucent-white, concealed in substrate, vertical from bulbil cluster, horizontal branches (stolons) potentially connecting the vertical flowering stems, stems unbranched, terete, 3–10.5 cm long (and possibly more), 1–2 mm diam., internodes 4.5–23(–40) mm long. Scale-leaves, alternate, translucent-white, slightly spreading, ovate or lanceolate, 2–4(–5.5) x 0.8–1.2 mm; axillary buds globose 0.5 mm diam. *Bulbil clusters* rare, or rarely preserved, underground, at base of stems, 4–7 cm below the flowers, (2–)8–12 bulbils per cluster, bulbils ellipsoid,, each 1.5–2 x 1–1.25 mm, each with an apical rootlet 5–7 mm long. *Flowers* developing from the stem apex in succession, 1–2 per stem, rarely 2 flowers open at a time per stem. Subtending bract translucent, forming a hood over the ovary and the dorsal surface of the lower perianth tube, concave, oblong or oblanceolate, 7–13 x 4 mm, apex rounded to acute. *Lower perianth tube* horizontal, shortly cylindrical to subellipsoid, 8–9 x 0.6–0.7 mm, c. 4.5 mm wide at base, translucent-white (sometimes with 6 longitudinal faint purple lines on the proximal part of the lower tube corresponding to the adnate filaments on the inner surface), outer and inner surface smooth, glabrous. *Annulus* or flange extending along the junction of the lower and upper perianth tubes, inserted c. 0.5 mm above the staminal filament insertion on the dorsal tube, absent on the ventral side, asymmetric, unlobed, glabrous and discontinuous, absent in the ventral 1/3 of the tube; in the dorsal 2/3 of the lower tube projecting at 90º into the tube cavity for up to 0.6 mm.

*Upper perianth tube* golden yellow-orange, paler at junction with lower tube, erect, forming an angle of 90º with the dorsal surface of the lower perianth tube, shortly cylindrical, c. 6 x 4 mm, apex rounded, the mouth horizontal, angled at 90–130º from the axis of the upper tube, surrounded by a corona, inner surface glabrous; outer surface of upper and lower perianth tube densely papillate (Fig. 3). Mouth orbicular or transversely elliptic, 5–7 mm diam., orbicular. Corona c. 0.5 mm long, projecting forward as a shortly cylindrical rim from the insertion of the tepals around the mouth. *Tepals* 6, ±monomorphic (subequal), spreading to ± forward-directed, white, at the base shaded yellow-brown, narrowly triangular-linear, (6.5–)7–10(–14) x 0.4–0.7 mm (1.5(–2) mm wide at base), not hastate, basal lobes absent, margins revolute at base, apex rounded, glabrous. *Stamens* 6, staminal filaments joined proximally to perianth tube for 2–3 mm, appearing as purple lines on the tube exterior surface, distally free, patent and arching downwards to the stigma, ±terete, 0.8–1(–1.6) x 0.2–0.3 mm; anthers angled downwards from filaments towards and apices attached to the stigma, oblong-elliptic in plan view, 1 x 0.5–0.7 mm, thecae 2, collateral; connective about 0.2 mm wide, leaving the anther cells visible dorsally; anther cells each 0.3 mm wide, adjacent, not separated by connective; distal connective appendage c 0.2 mm long, joined to upper surface of stigma. *Ovary* subturbinate, 2.5–3.5 x 4–4.5 mm; placentation not seen. Style cylindrical, c. 1 mm long, 0.1 mm diam. (dehydrated material), glabrous, stigma discoid, the centre concave dorsally, inconspicuously 6-lobed, c. 0.75 mm diam., lobes semicircular, c. 0.1 x 0.2 mm, erect. *Fruit globose*, c. 4 x 5 mm, only seen pre-dehiscence with perianth attached (immature). *Seeds* (from immature fruit) enclosed in a transparent pellicle, brown, irregularly narrowly ellipsoid, 0.5–0.7 x 0.2 mm, with longitudinal ridges, base (funicle) c. 0.08–0.15 x 0.08–0.12 mm; apex with dome (elaiosome?) 0.08–0.1 x 0.05 mm. Figs 1–4.

### DISTRIBUTION & HABITAT

Tanzania, Udzungwa Mts, Kihansi Gorge, in the lower part of the gorge in the area affected by the spray (then present) from the lower waterfall (see hydrology section below). Species-rich evergreen forest depending on the spray of the Kihansi falls (for further ecological data see Lovett et al. 1997 and discussion below); 770– 800 m elev.

### CONSERVATION STATUS

The forest habitat of *Afrothismia lovettii* has been negatively changed as a result of the construction of the Kihansi hydroelectric dam in 1999.

The dam diverted the Kihansi River into tunnels in the heart of the mountain where turbines produced more than 10% of Tanzania’s electricity when the scheme started operation in 2000. It also meant that the waterfalls no longer flowed in the dry season, substantially changing the moist microclimate of the gorge by removing the spray that kept the forest habitat of this species perhumid (see notes below). We consider it likely that this explains why neither this species, nor the other two microendemic mycoheterotrophic plant species *Kihansia lovettii* Cheek and *Kupea jonii* Cheek (collected at the same time by the second author), have been detected in the last 25 years since they were first discovered as part of the environmental impact studies for the dam by the second author in 1997 and the second collection in January 2001 by Peter Hawkes (pers. comm. to Cheek 2022). The extinction in the wild of the Kihansi toad *Nectophrynoides asperginis* Poynton, Howell, Clarke & Lovett, 1999, another micro endemic of the Kihansi falls, is well documented (e.g. Channing et al. 2006). It seems very likely that these three mycoheterotrophic plant species are also globally extinct. However, further searches should be made at the correct season in case they survive.

At the time of the type collection in March 1997, 100–200 individuals were noted at the only known site of about 50 m x 50 m (Lovett in Cheek 2004a), for which the altitude given was 770 m. The grid reference for *Lovett* 5063 was read off a map, and is now known to be erroneous as plotting on Google Earth Pro shows that it is outside the gorge at an elevation of only c. 300 m. In the same way, the grid reference given for the second collection was also misleading as it gives a site c. 40 km to the west in farmland. However, the altitude was given as 800 m, which is within the margin of error of the first, and so is possibly the same site, yet only three individuals were noted and collected (Peter Hawkes pers. comm. to Cheek 2022). However, exhaustive surveys for this species were not conducted at the time. Peter Hawkes is an entomologist and collected the *Afrothismia* for interest only. Using the required IUCN grid cells of 2 km x 2 km, gives an area of occupancy of 4 km^2^, and following IUCN guidance, the extent of occurrence can be taken as the same. Since only a single location is known, and habitat has been degraded (see above and below), our provisional IUCN assessment of *Afrothismia lovettii* is Critically Endangered (Possibly Extinct) (CR B1ab (iii)+2ab(iii)). We consider the species Possibly Extinct since it has not been seen in the last 25 years since it was recovered despite periodic site visits by biologists.

We recommend that specific surveys are made to attempt to rediscover *Afrothismia lovettii* in case it is not yet extinct. These should be conducted by specialists in heteromycotrophic plants, at the correct season for finding the plant i.e. in January to March. The site of collection was (JL pers. obs. 1997) a flat area of forest near the stream below the lower falls (mid gorge). On Google Earth Pro this equates to a polygon of c. 1.18 km^2^ or c. 9 ha bounded to the west by the river, the northern point being 8° 35′ 20” S, 35° 51′ 02”E and c. 855 m alt., the southern point 8° 35′ 28” S, 35° 50′ 55”E and c.748 m alt.

It is to be hoped that *Afrothismia lovettii* might yet be found elsewhere in the Udzungwa range. However, since the microclimate of the Kihansi falls gorge is unlikely to be replicated elsewhere (see studies below) the possibility of this seems low. Should the species be rediscovered, we recommend that a management plan is developed to ensure the survival of this species and is implemented following the protocols of Couch *et al*. (2022). This should include a public sensitisation and education programme, analysing and addressing the threats to the species, and annual monitoring of *Afrothismia lovettii* as well as the sympatric Critically Endangered sympatric *Kihansia lovettii* and *Kupea jonii* populations to determine trends in survivorship and threats, seedbanking. Cultivation and translocation of achlorophyllous mycotrophic flowering species such as *Afrothismia lovettii* has not yet been achieved but might be possible if sufficient resources and time can be allocated for researching this, for which the first step should be to determine the autotrophic partner species of the fungi upon which the plant depends. Without this information, planning for the translocation of any achlorophyllous mycotroph has a low chance of success.

### ADDITIONAL SPECIMEN EXAMINED

Tanzania, Kihansi Gorge, “8º 35’ S, 35º 31’ E” (erroneous), fl. Jan. 2001 P.G. Hawkes in *Ray Neyland* 1936 (recorded as 13 July 2001 in error) (LSU LSU00213308!)

### PHENOLOGY

Flower and fruit in January and March

### ETYMOLOGY

Named for Professor Jon Lovett, the collector of the type specimens. He is also commemorated in the names of the sympatric endemic species from Kihansi *Kupea jonii* and *Kihansia lovettii* (Triuridaceae, Cheek 2004a). *Salacia lovettii* N.Hallé & B.Matthew (Celastraceae, Hallé & Mathew 1986), *Trichilia lovettii* Cheek (Meliaceae, Cheek 1989) and *Psychotria lovettii* Borhidi & Verdc. (Rubiaceae, Borhidi & Verdcourt 1990), are examples of other species from Tanzania named in his honour.

### NOTES

The Hawkes in *Neyland* 1936 specimen at Louisiana State University herbarium (LSU) was discovered by the first author reviewing images of *Afrothismia* on gbif.org in 2022 and suspected to be new to science immediately. It was not connected to *Lovett* 5063 until later. The LSU specimen had originally been deposited at the herbarium of McNeese State University until that was amalgamated with LSU. It had preserved in alcohol but was rehydrated for DNA extraction and mounting on an herbarium sheet (notes on sheet).

The LSU herbarium specimen has the mouth of the perianth tube facing upwards, and the perianth lobes vertical, discordant from the posture in life and when spirit-preserved (Fig. 3). Some pressed individuals of *Lovett* 5063 also display this likely artefact of pressing.

## Discussion

### The Kihansi Gorge spray sustained forest: humidity studies

The Kihansi River of SW Tanzania is remarkable in that it has a fairly large catchment above the southeast facing escarpment of the Udzungwa mountains in contrast to other rivers flowing down the escarpment which have catchments more or less limited to the escarpment itself. In consequence the lower part of the Kihansi River maintains a relatively high year-round flow. This attribute, combined with the large elevation drop at the head of Kihansi Gorge, made the site suitable for the location of the Lower Kihansi Hydropower station. From a biological perspective, the sustained flow of the river, even in the dry season, and series of waterfalls in gorge, created a humid microclimate in the isolated forest patch that would have persisted for an exceptionally long period of time, for example throughout decreases in rainfall during the Pleistocene and probably earlier.

Between 1997 and 2001 a series of microclimatic measurements were made in Kihansi Gorge by James Taplin and Simon Tokumine as part of the Long-Term Environmental Monitoring Project to assess the impact of the Lower Kihansi Hydropower Project on the environment of Kihansi Gorge (NORPLAN, 2001). In 1997 temperature and humidity measurements were made during the period 10/9/97 - 27/9/97 (18 inclusive days) prior to diversion in 2000 of the Kihansi River to the hydropower turbines. The average of these data were used to create an interpolated map of gorge microclimate at different times of day. Figure 5 shows the humidity map at 14.00 hr, which is the hottest time of day. The colour purple is the maximum humidity and the colour red is the minimum.

**Fig. 5.**
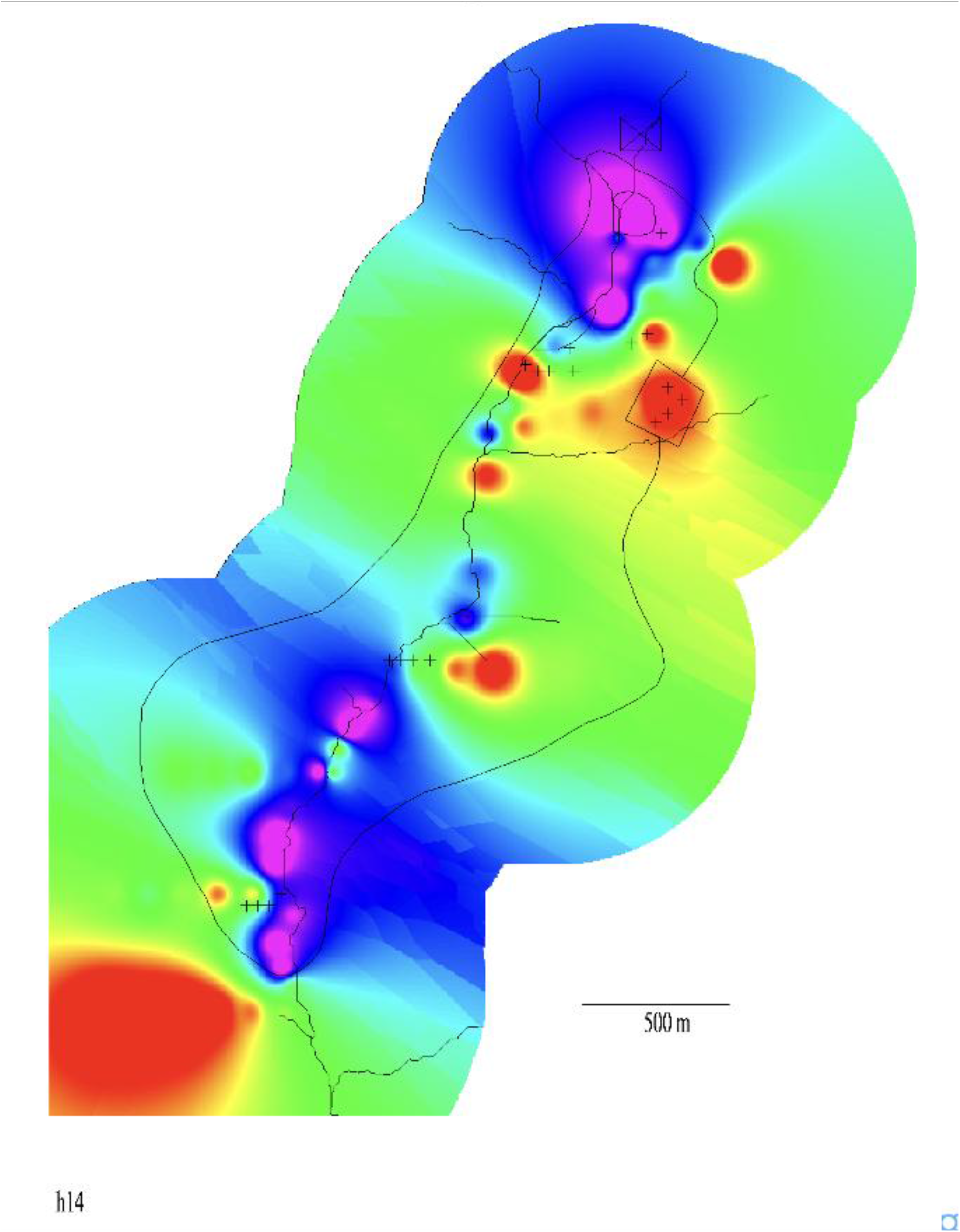
The Kihansi Gorge, Tanzania, showing humidity levels at 1400 hrs in September 1997, before the falls were diverted by the hydroproject. High humidity areas shown in blue and purple. Lowest humidity areas shown as red (from NORPLAN 2001).

The main Kihansi waterfall is in the upper part of the gorge (Figure 5 in 1997 prior to the river diversion) and generated significant quantities of spray to create an area of humidity represented by the upper purple and blue patch in Figure 5). There are secondary falls in the mid-gorge, which created another area of high humidity, again represented by the purple and blue areas on the map. The *Afrothismia* was collected from the area of high humidity in the mid-gorge forest.

### Kihansi Gorge Forest: the most important site for mycoheterotrophic plant species diversity

The coincidence of three species of mycoheterotroph, *Kihansia lovettii* and *Kupea jonii* (Triuridaceae), together with *Afrothismia lovettii*, makes (or made) the Kihansi gorge the most important site for species diversity of mycotrophic plants in eastern Africa. And in terms of numbers (three) of strict endemics of this lifeform, the Kihansi gorge is or was, the most important site in all Africa together with Mt Kupe in Cameroon which also has three strict endemic species of *Afrothismia* (Cheek et al. 2019; Cheek et al. 2004c), as well as slightly more widespread species such as *Kupea martinetugei* Cheek & S.A. Williams (Triuridaceae, Cheek et al. 2003). In contrast, the most species-diverse site in West Africa has only two site endemic species among the five species present (Cheek & van der Burgt 2010; Cheek et al. 2026). It is conceivable that additional species remain to be found, or once occurred at Kihansi, since the Udzungwa range in which Kihansi occurs supports or supported, other rare species of this life-form, notably *Afrothismia insignis* Cowley (Cowley 1988) and *Seychellaria africana* Vollesen (Triuridaceae, Vollesen 1982).

It is regrettable that the World Bank-funded hydroproject may have resulted in the global extinction of these three species of mycoheterotrophic plant. It is an example of the potential conflict between the need for development and conservation of biodiversity in sites where there are high numbers of narrow endemics

### Conservation importance and policy in the Eastern Arc and the Udzungwa Mts

The Udzungwa Mts of which the Kihansi Gorge is part, are among the mountain and coastal forests of Tanzania which have long been recognised as an area of botanical interest (Polhill 1968). A map of the Eastern Arc mountains delimiting the area of high endemism was published in 1985 (Lovett 1985) and an estimate of the floristic endemism in the forests of Tanzania and Southeastern Kenya was presented at the 1985 AETFAT Symposium at the Missouri Botanical Gardens in St Louis, USA (Lovett 1988). Botanical exploration during the early 1980s revealed many new species (Lovett *et al*. 1988) and in consequence the Eastern Arc were identified as a ‘biodiversity hotspot’ by Norman Myers in 1990 (Myers 1990) due to a combination of the high numbers of restricted range species; habitat fragmentation; and ongoing threats from logging and conversion to agriculture leading to forest loss. Designation as a globally important biodiversity hotspot led to increased concern for conservation and funding for conservation activities during the 1990s supported by changes in forest policy and law (Lovett 1992, 2003a, b; Lovett & Pócs 1993). For example, the Tanzanian government established the Eastern Arc Mountains Conservation Endowment Fund as a Not-for-Profit Non-Governmental Organization (NGO) in 2001 as a means of channelling resources from international donors to field-based activities. Conservation activity is continuing to grow and new species are still being discovered, as evidenced in this paper.

### New discoveries of plant species in the Udzungwa mountain forests of Tanzania

The discovery of *Afrothismia lovettii* is only the latest endemic new species to science to be discovered from the Udzungwa Mts, one of the well-known Eastern Arc Mountains of Tanzania. Apart from the other endemic heteromycotrophs already discussed above, among the other sp. novs described since the turn of the century are *Pauridiantha udzungwaensis* Ntore & Dessein (Ntore et al. 2003), *Ancistrocladus tanzaniensis* Cheek (Ancistrocladcaeae, Cheek et al. 2000; Cheek 2000), *Lukea triciae* Cheek (Cheek et al. 2022), *Vepris lukei* and *V. udzungwa* Cheek (Rutaceae, Cheek & Luke 2023) and most recently *Begonia wahehe* A.Bianchi, L.Angelini & Q.Luke (Begoniaceae, Bianchi et al. 2026).

## Conclusion

Until new species such as *Afrothismia lovettii* are formally named, they are essentially invisible to science, and Red Listing, which can be effective in improving conservation outcomes (Cheek et al. 2020b) is difficult. Since three out of four plant species have been found to be threatened at the point of publication currently (Brown et al. 2023), and destruction and degradation of natural habitat continues apace, including from infrastructure projects such as the Kihansi dam, it is urgent to find the remaining unknown plant species on the planet so that they can also be named, and have the possibility of being protected from extinction, for example by being included in Important Plant Areas (Darbyshire et al. 2017). However, the lesson of this paper seems to be that even when environmental impact studies are carried out in advance of major projects, and highly threatened species are found to be present, they may still become extinct. It appears that species in several genera of non-photosynthetic heteromycotrophs are especially at risk of global extinction due to their very small range sizes and specialised and poorly understood micro-habitat requirements, with extinction already recorded reliably elsewhere in tropical Africa (e.g. Cheek & Onana 2011; Cheek et al. 2018; 2019).

## Acknowledgements

The staff at the herbarium of Louisiana State University (LSU), especially Collections Manager Jennifer S Kluse are thanked for their excellent efforts in facilitating access to the material of this new species to science that they hold.

Peter Hawkes, entomologist at Afribugs (www.afribugs.com) is thanked for dialogue on his collection of the species at Kihansi Gorge and for his excellent photos of the spirit collection that he preserved. We thank two anonymous reviewers for constructive comments on an earlier version of this manuscript.

## Declaration

The authors declare they have no conflict of interest.

